# A stealth adhesion factor contributes to *Vibrio vulnificus* pathogenicity: Flp pili play roles in host invasion, survival in the blood stream and resistance to complement activation

**DOI:** 10.1101/613810

**Authors:** Tra–My Duong–Nu, Kwangjoon Jeong, Soo Young Kim, Wenzhi Tan, Sao Puth, Kwang Ho Lee, Shee Eun Lee, Joon Haeng Rhee

## Abstract

The tad operons encode the machinery required for adhesive Flp (fimbrial low-molecular-weight protein) pili biogenesis. *Vibrio vulnificus*, an opportunistic pathogen, harbors three distinct *tad* loci. Among them, only *tad1* locus was highly upregulated in *in vivo* growing bacteria compared to *in vitro* culture condition. To understand the pathogenic roles of the three *tad* loci during infection, we constructed single, double and triple tad loci deletion mutants. Interestingly, only the *Δtad123* triple mutant cells exhibited significantly decreased lethality in mice. Ultrastructural observations revealed short, thin filamentous projections disappeared on the *Δtad123* mutant cells. Since the pilin was paradoxically non-immunogenic, a V5 tag was fused to Flp to visualize the pilin protein by using immunogold EM and immunofluorescence microscopy. The *Δtad123* mutant cells showed attenuated host cell adhesion, delayed RtxA1 exotoxin secretion and subsequently impaired translocation across the intestinal epithelium compared to wild type, which could be partially complemented with each wild type operon. The *Δtad123* mutant was susceptible to complement-mediated bacteriolysis, predominantly via the alternative pathway, suggesting stealth hiding role of the Tad pili. Taken together, all three *tad* loci cooperate to confer successful invasion of *V. vulnificus* into deeper tissue and evasion from host defense mechanisms, ultimately resulting in septicemia.

**Author Summary:** To understand the roles of the three Tad operons in the pathogenesis of *V. vulnificus* infection, we constructed mutant strain with single, double and triple Tad loci deletions. Employing a variety of mouse infection models coupled with molecular genetic analyses, we demonstrate here that all three Tad operons are required for *V. vulnificus* pathogenicity as the cryptic pili contribute to host cell and tissue invasion, survival in the blood, and resistance to complement activation.

## Introduction

*Vibrio vulnificus* is an opportunistic marine pathogen that causes fatal septicemia and necrotizing wound infections in susceptible individuals with underlying hepatic diseases and other immunocompromised conditions. In humans, this pathogen frequently causes rapidly progressing fatal sepsis with a mortality rate of greater than 50% within a few days post-infection (1-4). During the infectious process, *V. vulnificus* must cope with dramatic environmental changes by sensing changes in the host milieu (5). To establish successful infections *in vivo, V. vulnificus* must manage spatiotemporally coordinated changes in the expression levels of various virulence genes.

To understand the genome-wide gene expression changes in *V. vulnificus* after infection, we recently performed a transcriptomic analysis of cells grown *in vivo* using a rat peritoneal infection model. Notably, among the newly identified *in vivo*-expressed genes, a Flp/Tad pilus-encoding gene cluster (the *tad1* locus) was found to be highly upregulated *in vivo* (unpublished data). Flp pili are polymers of the mature Flp pilin protein, and they are assembled and secreted by a complex of proteins encoded by the *tad* operon. Flp pili were reported to be abundantly expressed, extremely adhesive, and bundled in *Aggregatibacter* (previously *Actinobacillus*) (6-9). The Tad proteins have been reported to be essential for adherence, biofilm formation, colonization, and pathogenesis in a number of genera and are considered to be instrumental in the colonization of diverse environmental niches (6, 7, 10-12).

*V. vulnificus* CMCP6 harbors three distinct *tad* loci (13) (S1 Fig), among which the *tad1* locus has been identified as a possible virulence factor because of its ubiquity in sequenced virulent *V. vulnificus* strains (14-16). In contrast, the *tad3* locus was expressed at a higher level in artificial seawater than that in human serum (17). This study attempted to investigate the contribution of the high *in vivo* expression of the *tad1* operon to the *V. vulnificus* pathogenicity and to understand why three similar *tad* operons were maintained throughout the long history of evolution. We evaluated how each *tad* operon contributes to *V. vulnificus* virulence. Since the three *tad* operons share genes with similar function, single-gene-mutation analyses could not rule out overlapping functions of the remaining genes in the same operon or in the other *tad* operons. Thus, we constructed mutant strains with single and multiple complete *tad* loci deletions and then complemented them with individual cosmid clones harboring each *tad* operon. Using a variety of mouse infection models coupled with molecular genetic analyses, we demonstrate here that all three *tad* operons are required for *V. vulnificus* pathogenicity as the cryptic pili contribute to host cell and tissue invasion, survival in the blood, and resistance to complement activation.

## Results

### Transcriptional analyses of the three structural *flp* genes in *V. vulnificus* infecting the rat peritoneal cavity

To understand how host signals modulate *tad* operon expression in *V. vulnificus*, we analyzed the *in vivo* transcriptional levels of three structural *flp* genes using a rat peritoneal infection model. Real time RT-PCR results indicated a significantly higher *flp-1* mRNA level when the bacteria were grown *in vivo*, corresponding to an approximately 878-fold increase (Fig. 1A) (*P <* 0.001). Conversely, both the *flp-2* and *flp-3* transcript levels were slightly decreased when *V. vulnificus* was grown *in vivo* (Fig. 1A) (*P <* 0.05 for *flp-2* and *P <* 0.001 for *flp-3*). The expression levels of the *flp* genes were also measured using conventional RT-PCR. Using different numbers of amplification cycles, we confirmed that the *flp-2* and *flp-3* genes were transcribed at low levels under both tested conditions; in particular, *flp-2* expression was detected only after 35 cycles (Fig. 1B).

**Figure 1.**
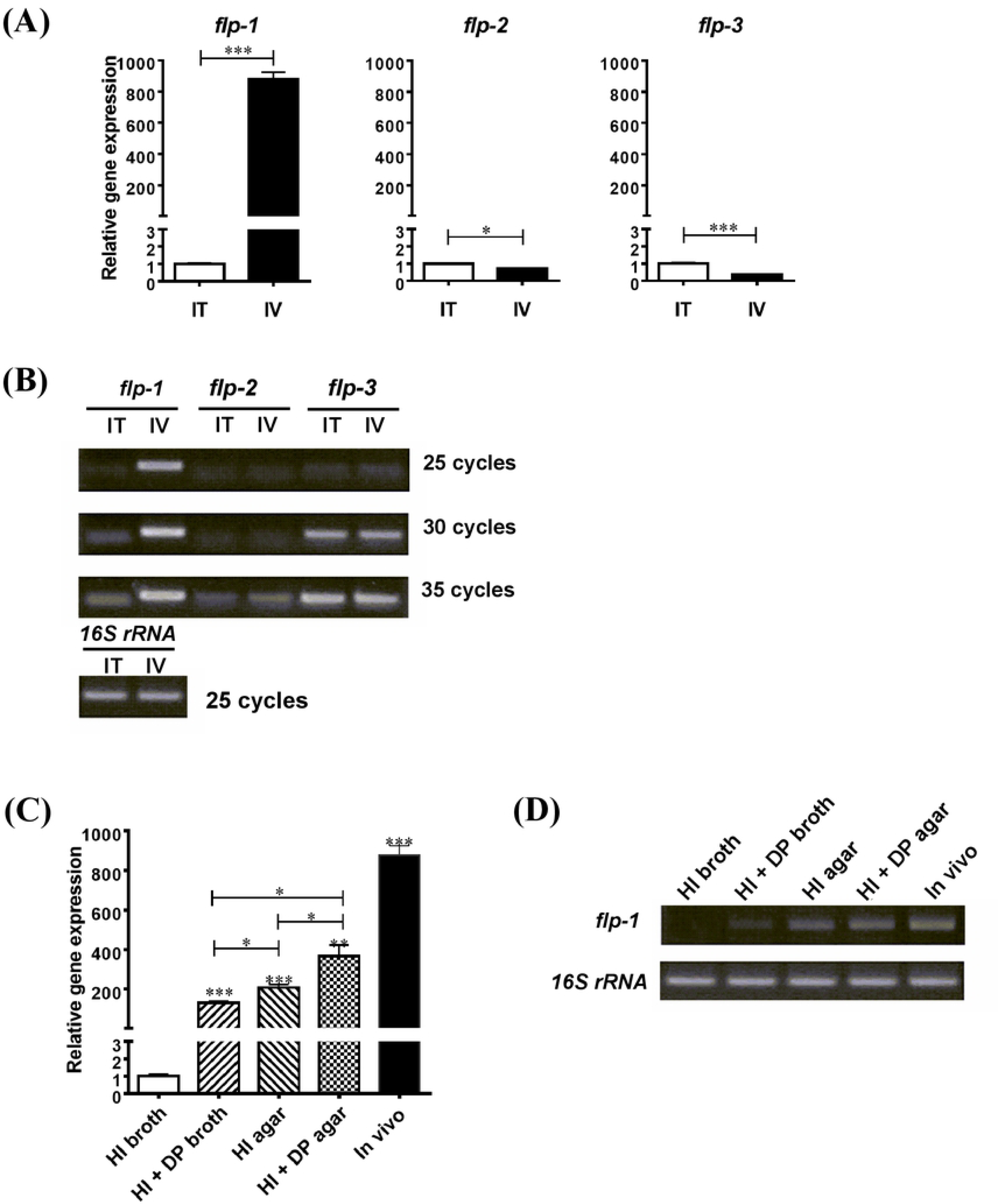
Transcriptional analyses of structural *flp* genes in three *tad* operons of *V. vulnificus*. RNA was isolated from bacteria grown in the rat peritoneal cavity (IV, *in vivo*) or in 2.5% NaCl HI broth (IT, *in vitro*) and the transcript levels of the structural *flp* genes were analyzed via real-time **(A)** and conventional RT-PCR **(B)**. RNA was isolated from bacteria grown under 2.5% NaCl HI broth, iron-limited, solid surface and in vivo conditions and the transcript levels of the structural *flp* genes were analyzed via real-time **(C)** and conventional RT-PCR **(D)**. DP (80 µM) was added to 2.5% NaCl HI broth for iron limitation. The real-time RT-PCR data were normalized to *gyrA* and expression relative to the *in vitro* level. Data shown represent the mean ± SEM of three independent experiments performed in triplicate. Statistical analysis was carried out using Student’s *t* test (*, *P <* 0.05; **, *P <* 0.01; ***, *P <* 0.001).

In a wide variety of bacteria, type IV pili expression is solid-surface dependent (18, 19), and the *tad1* locus was recently found to be expressed under iron-limited conditions (13). Thus, we measured the expression of the *flp-1* gene under these growth conditions. As shown in Fig. 1C and D, both the iron-limited and surface-associated growth conditions clearly stimulated *flp-1* transcription, increasing its expression levels by approximately 131- and 210-fold, respectively (*P <* 0.001 compared to that of the expression level in HI broth). Combining these two conditions significantly increased the *flp-1* transcript level by 367-fold (*P <* 0.01), but the transcription level was still much lower than that observed *in vivo*. This finding indicates that changing one or two growth parameters in culture does not mirror the *in vivo* environment, where multiple host factors and growth conditions would simultaneously influence *tad1* operon expression.

### All three *tad* operons are required for full *V. vulnificus* virulence

To explore the contribution of each *tad* operon to *V. vulnificus* pathogenicity, we performed mouse lethality assays employing intraperitoneal (i.p.) and intragastric (i.g.) infection routes. Interestingly, in the i.p. infection model, the Δ*tad123* mutant showed a 41-fold increase in the LD_50_, while the single and double mutants showed no differences (Table 1). Significantly prolonged survival was observed in the Δ*tad123* mutant-administered mice, which received infectious doses of 1.0 × 10^7^ and 1.0 × 10^6^ CFU/mouse. At a dose of 10^7^ CFU/mouse, all of the mice infected with wild-type cells died within 5 hours post-infection, whereas approximately 60% of the mice infected with the Δ*tad123* mutant survived up to 48 h after the challenge (S2 Fig) (*P <* 0.01). However, after i.g. infection, which leads to slower translocation of the bacteria into blood circulation, we observed only a 10-fold LD_50_ increase (Table 1). The lethality varied depending on the route of infection, which influences the rate of bacterial invasion, growth and/or clearance at both the primary infection site and in the blood stream. Taken together, all three *tad* operons must be deleted to significantly ameliorate *V. vulnificus* virulence.

**Table 1.** Effect of the mutation of *tad* operons on the lethality for mice.

| *V. vulnificus*<br>strains | LD<sub>50</sub> (95% confidence limits) |  |
| --- | --- | --- |
|  | Intraperitoneal infection | Intragastric infection |
| WT | $5.5 \times 10^5$ | $1.1 \times 10^5$ |
| $\Delta tad123$ | $2.3 \times 10^7$ | $5.5 \times 10^6$ |
| $\Delta tad1$ | $2.1 \times 10^5$ | |
| $\Delta tad2$ | $5.5 \times 10^5$ | |
| $\Delta tad3$ | $4.0 \times 10^5$ | |
| $\Delta tad12$ | $5.5 \times 10^5$ | |
| $\Delta tad13$ | $7.0 \times 10^5$ | |
| $\Delta tad23$ | $6.4 \times 10^5$ | |
| Fold increase<br>(WT vs $\Delta tad123$) | 41 | 10 |

### The Δ*tad123* mutant exhibits significantly decreased adhesion to HeLa cells

Since common pili are generally involved in the attachment of bacteria to surfaces in nature, we hypothesized that deletion of the three *tad* operons might influence the adhesive ability of *V. vulnificus*. To test this hypothesis, we performed an adhesion assay in which HeLa cells were infected with *V. vulnificus* at an MOI of 250 followed by quantification of the number of bacteria adhered to the host cells. After incubation for 45 min, the number of Δ*tad123* mutant cells adhered to the HeLa cells was 13-fold less than that of the parental wild-type strain (Fig. 2A) (*P <* 0.001). The wild-type strain formed small clusters of aggregated bacteria on the surfaces of the HeLa cells, eventually leading to cell lysis. In contrast, only a few Δ*tad123* mutant cells attached to the surfaces of the HeLa cells, and the infected host cells maintained cell contours similar to those of the uninfected cells. However, the adhesion of Δ*tad123* mutant cells to the host cells gradually increased in a time-dependent manner (Fig. 2B, *P* < 0.01). Complementation with the *tad1* or *tad3* operon (S3 Fig), but not the *tad2* operon, significantly rescued the adhesive ability of the Δ*tad123* mutant cells (Fig. 2A) (*P* < 0.001 for *tad1* and *tad3, P* > 0.05 for *tad2*).

**Figure 2.**
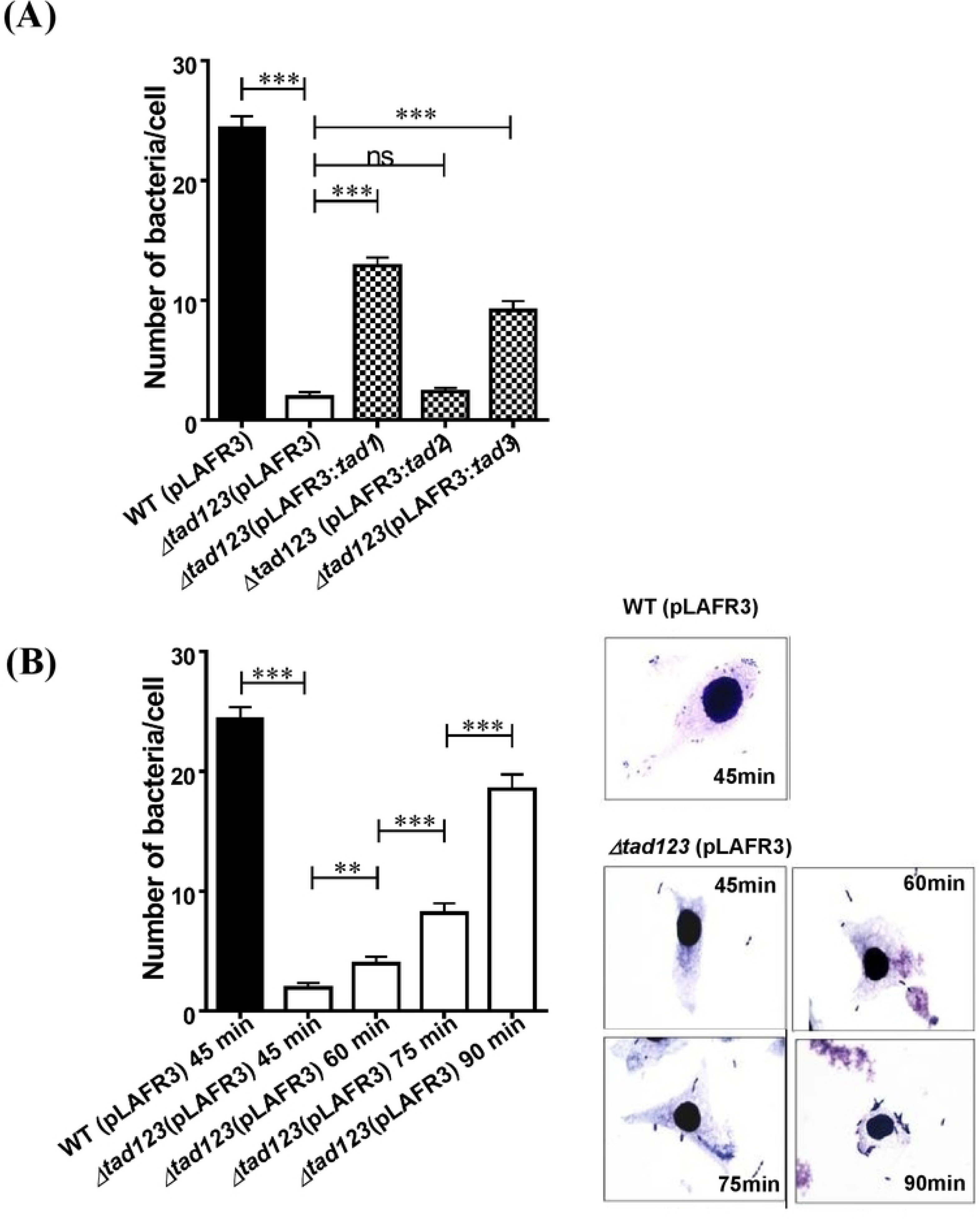
Significantly decreased adhesion to host cells by the Δ*tad123* mutant (A) and time-dependent recovery (B). HeLa cells were treated with log-phase *V. vulnificus* cells at an MOI of 250, and the bacterial cells that adhered to single HeLa cells were counted at the indicated time points. The morphology of the infected HeLa cells was observed after Giemsa staining at ×1,000 magnification. Data shown represent the mean ± SEM of five independent experiments performed with 30 replicates. Statistical analysis was carried out using Student’s *t* test (***, *P* < 0.001; ns, not significant compared to the wild-type strain).

### Structure of the Tad pili on the surface of *V. vulnificus* cells

To observe the morphology of the Tad pili, we prepared *in vivo* grown wild-type, Δ*tad123*, and Δ*tad123* cells carrying pLAFR3::*tad1* locus then performed scanning electron microscopy (SEM) observation (Fig. 3). In the wild-type strain, the cell surface appeared to be covered with slime-like material. Corrugated elevations and grooves ran along the longitudinal axis. In the Δ*tad123* mutant strain, which was devoid of the slime-like materials, the grooves and elevations were more conspicuous compared with those of the isogenic wild-type strain. Moreover, the directionality of the convexity of the surface structure was absent on the mutant surface. Interestingly, the cell surfaces of the *in trans tad1* complemented strains showed similar structural characteristics to the wild type strain, suggesting that the Tad pili contribute to the formation of the slime-like surface structure. However, the typical surface groove and convexity was less obvious in the complemented strains. These results suggest that the Tad pili of *V. vulnificus* might also contribute to the cell envelops biogenesis by including the slime-like outer structure.

**Figure 3.**
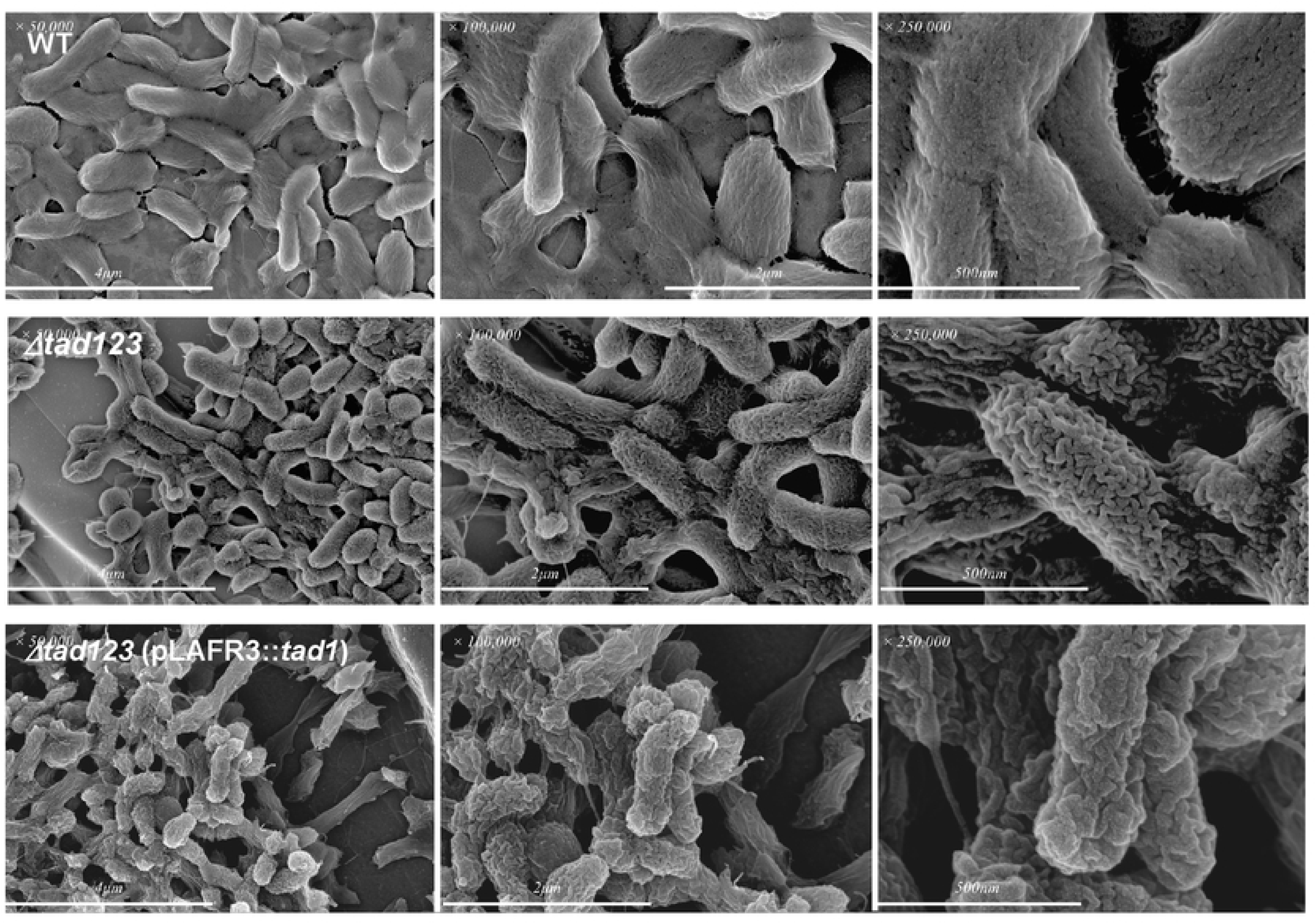
Ultrastructural observation of *V. vulnificus.* Bacteria were grown *in vivo* using a rat peritoneal infection model. The specimens were examined by SEM.

We tried to further characterize the thin fimbrial projections of the putative Flp pili via immunogold electron microscopy. Firstly, we produced recombinant Flp pilin proteins and attempted to raise specific antibodies against them in animals. However, we could not obtain any appreciably immunogenic antisera even after many repeated trials. Peptide-based immunizations were also unsuccessful in raising specific antibody responses. We became to conclude that the Flp pilin have very low immunogenicity. To solve this problem, we constructed *V. vulnificus* strains carrying a pBAD24::FlpV5 plasmid expressing a hybrid protein of Flp pilin fused to the highly immunogenic V5 tag (S4 Fig) with the expectation that the plasmid-encoded FlpV5 subunits would assemble into growing pilus fibers under the control of an arabinose-inducible promoter. By using dot blot analysis with an anti-V5 antibody, we confirmed the expression and assembly of FlpV5 into authentic pili in plasmid-harboring *V. vulnificus* transconjugants under the inducing conditions. We specifically detected positive signals in wild-type *V. vulnificus* cells (Fig. 4A), while no signal was detected in the Δ*tad123* mutant cells. The immunogold-labeling analysis revealed that only a small subpopulation of wild-type cells displayed gold-labeled Flp filaments on their surfaces, suggesting competition between the natural Flp and FlpV5 pilin subunits (Fig. 4B). Supporting the results of the immunogold staining, immunofluorescence detection via confocal microscopy also revealed positive fluorescent signals for V5p in the FlpV5-expressing wild-type strain (Fig. 4C). In contrast, the Δ*tad123* mutant cells did not show any fluorescent signals, and this deficiency was complemented *in trans* by cosmids encoding the wild-type *tad1* locus (Fig. 4C). Many broken filaments were found in the backgrounds of the EM photos, suggesting detachment of the brittle pilus structures during the sample preparation procedures. Taken together, the EM analyses demonstrated the obvious existence of extracellular Flp structures in wild-type *V. vulnificus* CMCP6.

**Figure 4.**
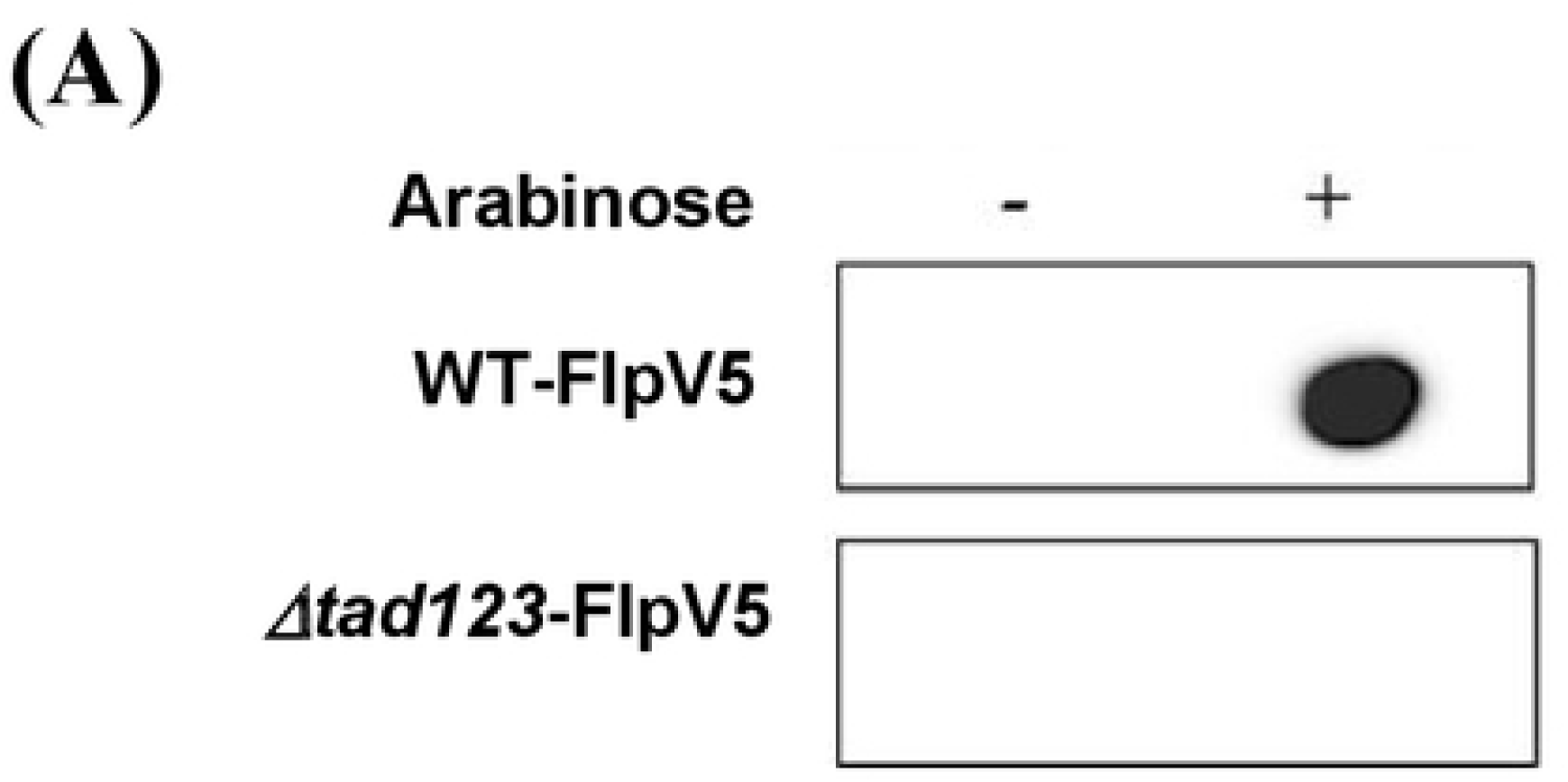
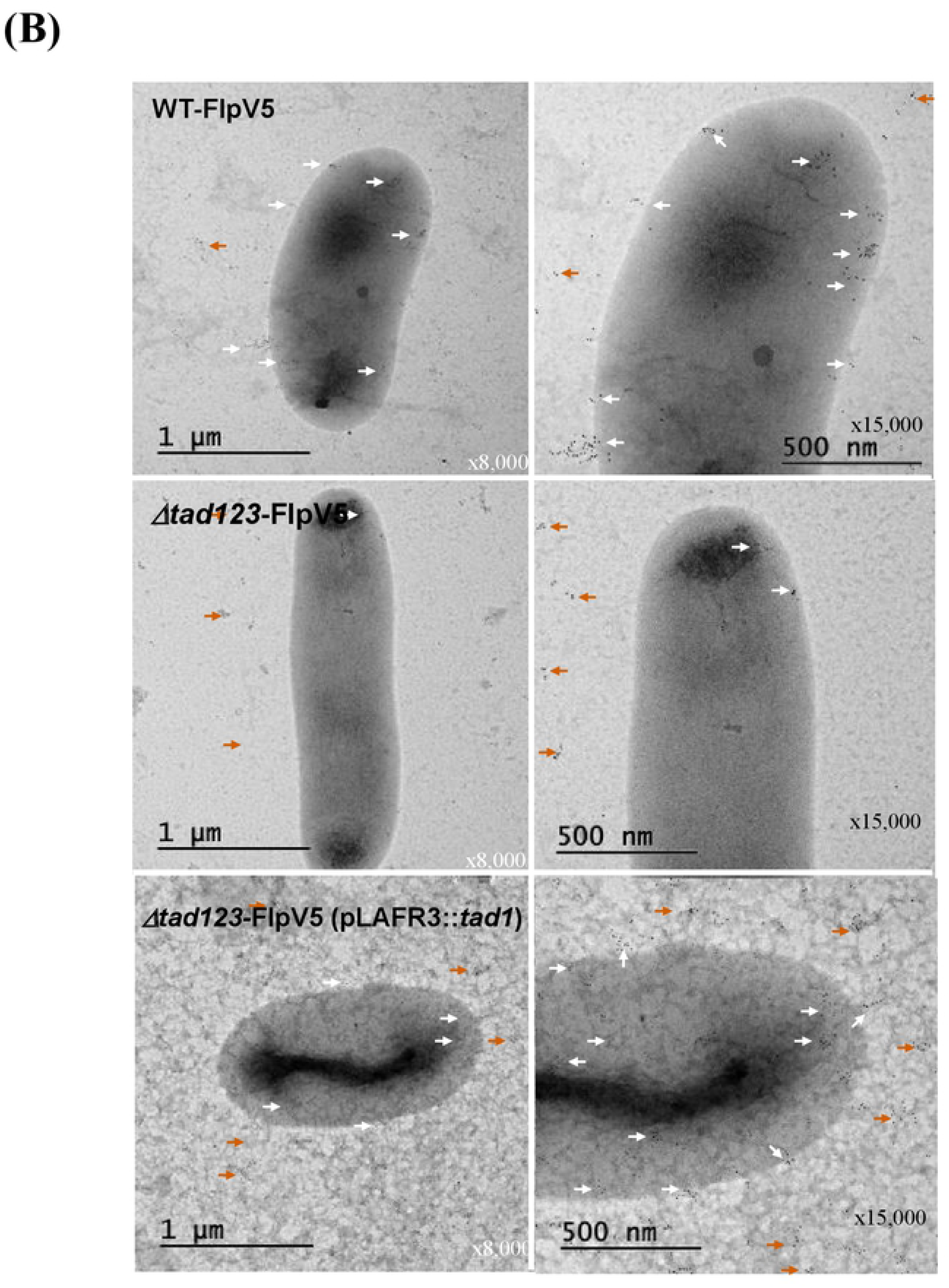
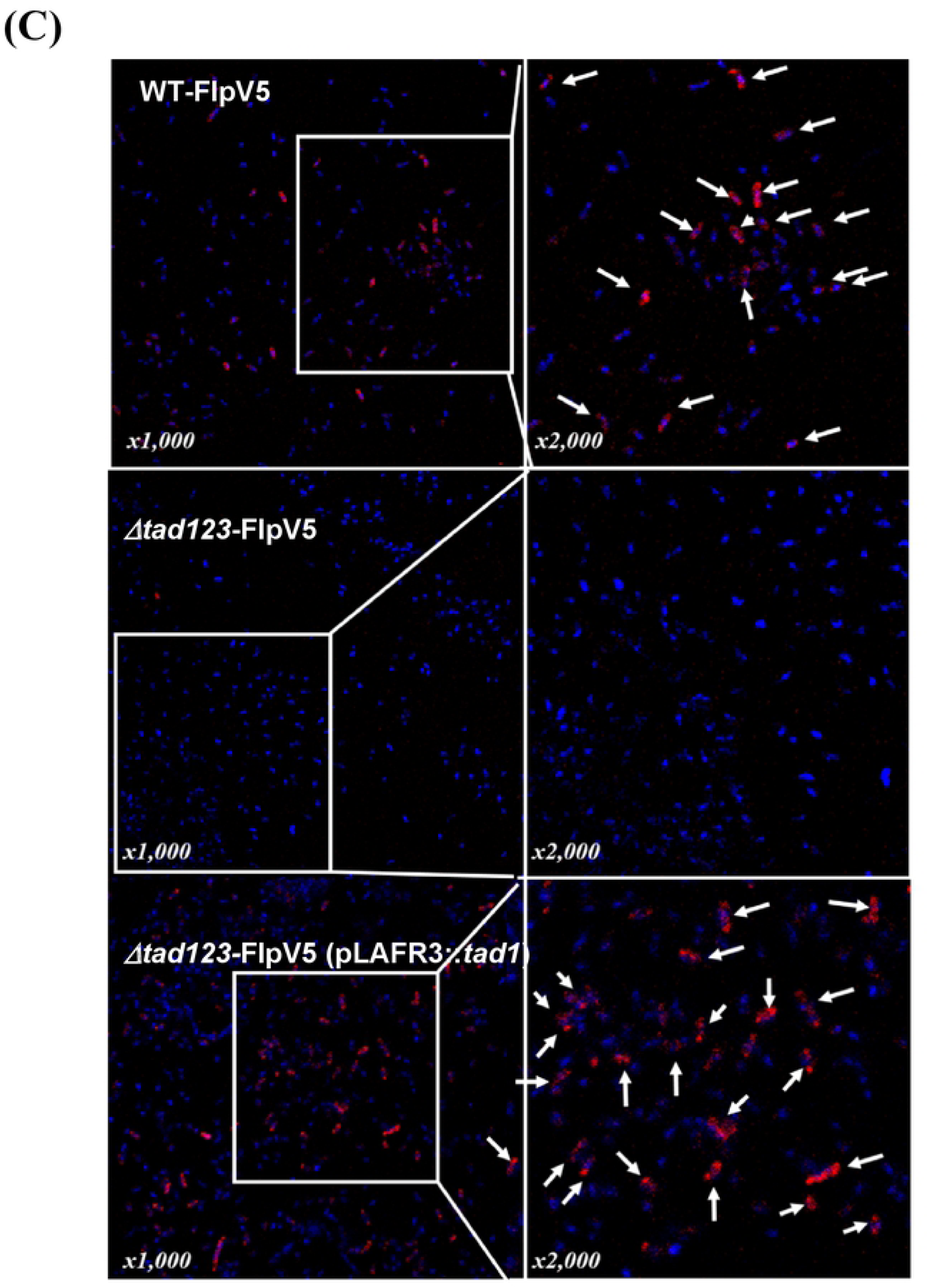
Detection of V5-tagged Flp. (A)Dot blot analysis. Bacteria were grown on a 2.5% NaCl HI-Amp agar plate supplemented with (induced) or without (non-induced) 0.1% arabinose for 4 h. The Flp-V5 fusion protein was detected using an anti-V5 polyclonal antibody. Δ*tad*-FlpV5 indicates the Δ*tad123* mutant strain carrying pBAD24 expressing the Flp-V5 fusion protein, and WT-FlpV5 indicates the wild-type strain carrying pBAD24 expressing the Flp-V5 fusion protein. **(B)** I**mmunogold labeling and TEM analysis.** Bacteria were fixed and incubated with an anti-V5 polyclonal antibody and a 5-nm colloidal gold conjugated goat anti-rabbit IgG secondary antibody. Only *V. vulnificus* cells expressing the Flp-V5 fusion protein showed positivity for immunogold particles on the cell surface. **(C) Immunofluorescent detection of Flp-V5**. Bacterial cells were visualized by staining their DNA with DAPI (blue), and anti-V5 detection appears in red (white arrowheads).

### The Δ*tad123* mutant cells display defective RtxA1 production, leading to delayed cytotoxicity toward HeLa cells

RtxA1 is a crucial cytotoxin involved in cellular damage and necrosis of infected tissues (20-24). We previously reported that host cell contact is required for RtxA1 production and cytotoxicity (20). Thus, we speculated that attenuated adherence to host cells should hamper RtxA1 production and consequently attenuate host cell killing and tissue invasion. We performed a Western blot analysis to assess RtxA1 production after HeLa cell infection. The toxin was detected using an anti-GD domain antibody targeting the C-terminal fragment (RtxA1-C; approximately 130 kDa), which is internalized in the host cell cytoplasm (25). As a result of its impaired ability to maintain contact with its host cells, the Δ*tad123* mutant exhibited significantly lower toxin production compared with that of its parental strain (Fig. 5A). RtxA1 was secreted in a time-delayed manner in the mutant cells, and its secretion gradually increased over time. This delay was significantly rescued by *tad* operon complementation, except in the case of the *tad2* (Fig. 5A). We next assessed the cytotoxicity of *V. vulnificus* toward HeLa cells over a time course. As shown in Fig. 5B, the Δ*tad123* mutant cells showed significantly delayed cytotoxicity toward HeLa cells (*P* < 0.001), whereas the single and double mutants showed no changes (*P* > 0.05). The cytotoxicity of the Δ*tad123* mutant cells approached the wild-type level after 2.5 h of incubation. This delay in the manifestation of the cytotoxicity was significantly recovered via *in trans* complementation with either the *tad1* or the *tad3* operons (Fig. 5C) (*P* < 0.01 for *tad1* and *P* < 0.001 for *tad3*). To investigate the possibility that this result might have been due to bacterial growth retardation in the HeLa cell culture medium, we examined the growth profiles of test strains in high-glucose Dulbecco’s Modified Eagle’s Medium (DMEM) (S5 Fig). No growth difference was observed between the wild-type and Δ*tad123* mutant strains. These findings, together with the LD_50_ results, highlight the significance of the three *tad* operons for the adhesion-mediated virulence of *V. vulnificus*.

**Figure 5.**
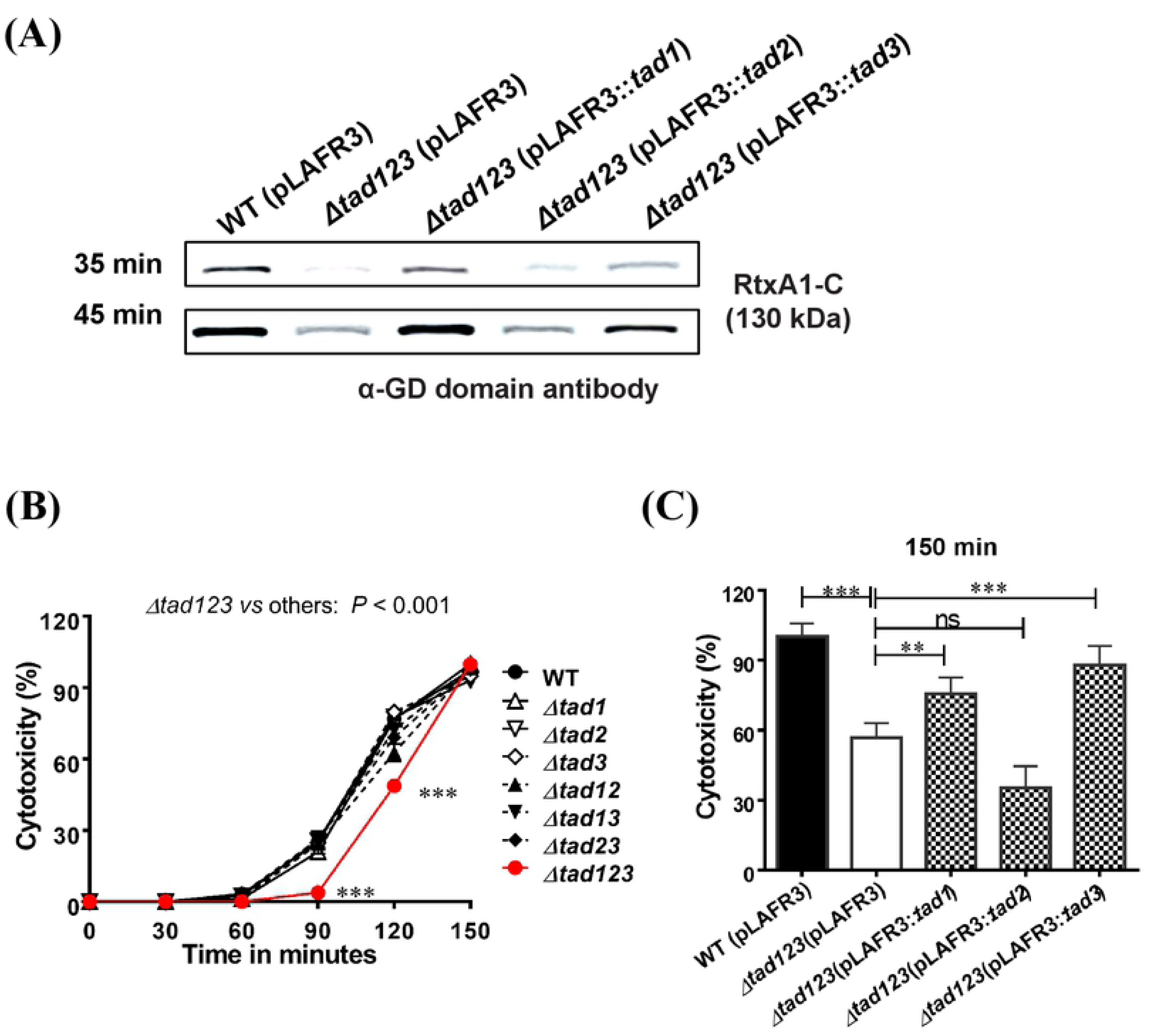
Delayed RtxA1 secretion (A) and cytotoxicity (B) by the Δ*tad123* mutant cells. **(A)** For RtxA1 detection, log-phase *V. vulnificus* cells were incubated with HeLa cells in 6-well plates at an MOI of 100 for 35 or 45 min. The cells in each well were lysed with lysis buffer, followed by concentration using the Amicon Ultra-0.5 centrifugal filter apparatus. The RtxA1 toxin was detected by Western blot analysis using an anti-GD domain antibody (RtxA1-C, a band of approximately 130 kDa). **(B)** Effect of *tad* operon mutations on cytotoxicity against HeLa cells. **(C)** Restoration of cytotoxicity in *tad*-complemented strains at 2.5 h post-incubation. Data shown represent the mean ± SEM of three independent experiments performed with five replicates. Statistical analysis was carried out using two-way ANOVA followed by the Bonferroni *post hoc* test (a) or Student’s *t* test (b). **, *P* < 0.01; ***, *P* < 0.001; ns, not significant.

### Tad pili are essential for intestinal invasion by *V. vulnificus*

Bacterial pili are used to attach to host cells and tissues, and confer invasive competence (26-30). Furthermore, secretion of the RtxA1 cytotoxin, which is induced by adhesion of the bacterial cells to the host cells, is highly correlated with host tissue invasion (20). To investigate the effects of mutation of the *tad123* loci on *V. vulnificus* invasion, we carried out an *in vivo* invasion assay using a mouse ligated ileal loop infection model. The viable bacterial cells in the blood of the infected mice were quantified to evaluate tissue invasiveness of bacteria. The number of bacterial cells in the blood samples from the Δ*tad123* mutant-infected mice was significantly lower than that in the mice infected with the wild-type strain, even 6 hours post-infection (Fig. 6A) (*P* < 0.001).

**Figure 6.**
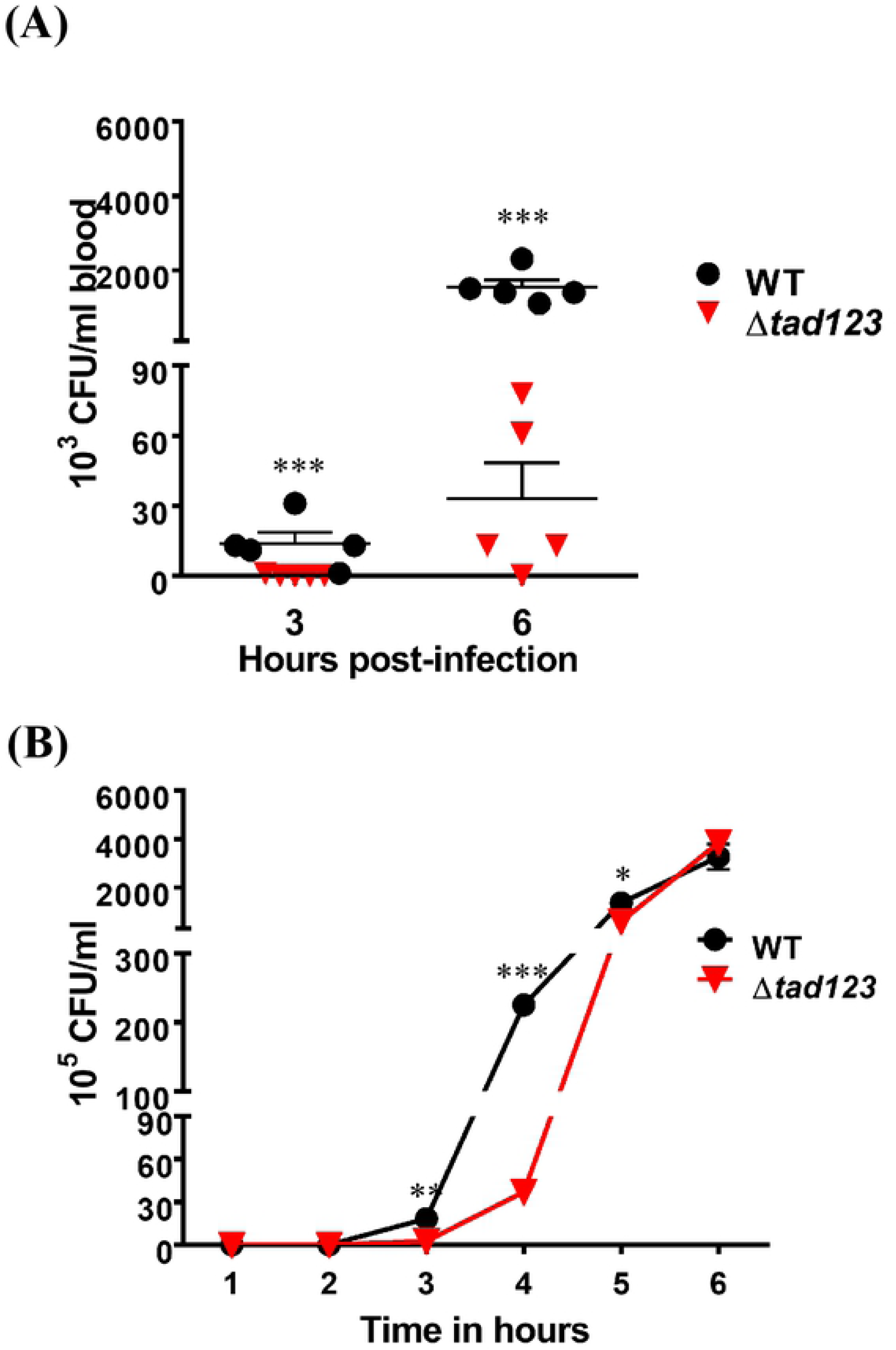
Effect of the triple *tad* operon mutation on *in vivo* and *in vitro* invasion. **(A)** *In vivo* invasion. Log-phase bacteria were inoculated into ligated ileal loops of mice. Blood samples were acquired from infected mice by cardiac puncture at the indicated times. Viable cells were counted by plating on 2.5% NaCl HI agar plates. **(B) *In vitro* invasion.** HCA-7 cells grown on Transwell filters were apically exposed to log-phase bacteria. Invasiveness was determined by measuring the number of bacterial cells that translocated from the apical to the basolateral compartment of the Transwells. Viable bacterial cells were counted by plating on 2.5% NaCl HI agar plates. Data shown represent the mean ± SEM of three independent experiments performed in five mice. Statistical analysis was carried out using Student’s *t* test (A) or two-way ANOVA followed by the Bonferroni *post hoc* test (B). *, *P* < 0.05; **, *P* < 0.01; ***, *P* < 0.001; ns, not significant. WT, wild-type *V. vulnificus*.

In addition to the *in vivo* invasion assay, bacterial invasiveness was further confirmed using an *in vitro* intestinal epithelial barrier system. Polarized HCA-7 cells grown on Transwells were apically infected with bacteria, leading to physical apical-to-basolateral trans-epithelial migration of the bacteria. After 3 to 5 hours of incubation, we detected significantly fewer Δ*tad123* mutant cells than wild-type cells in the basolateral chamber (Fig. 6B) (*P <* 0.05). The cell count of the Δ*tad123* mutant reached that of the wild-type strain after 6 hours of incubation. Taken together, the *in vitro* data and the *in vivo* invasion results demonstrate a function for Tad pili in conferring invasive competence to *V. vulnificus*.

### Tad pili are important for *V. vulnificus* survival in blood

It is likely that the impaired invasion alone could not fully account for the higher LD50 observed for the Δ*tad123* mutant in the intraperitoneal infection model (Table 1); therefore we hypothesized that mutation of the *tad123* loci could compromise *V. vulnificus* survival in the bloodstream. To investigate this possibility, we monitored the number of viable bacteria in the blood over a time course following intraperitoneal (i.p.) and intravenous (i.v.) infection. Interestingly, the triple mutant cells were defective at surviving in mouse blood. Significantly fewer mutant cells were recovered from the blood of mice infected via both routes (Fig. 7A and 7B). In particular, very few mutant cells were detected after direct introduction of the bacteria into the blood stream via i.v. injection, resulting in an approximately 3-log reduction in the number of CFUs compared with the number of CFUs detected for the wild-type strain (Fig. 7A). The viable bacteria in the blood following i.p. infection should represent *V. vulnificus* cells that succeeded at both invasive translocation and resisting the serum bactericidal activities. On the other hand, the i.v. infection model shows how well the wild-type and mutant bacteria survived the serum bactericidal activities. These findings clearly indicate that Tad pili also play important roles in the survival of *V. vulnificus* in the blood stream.

**Figure 7.**
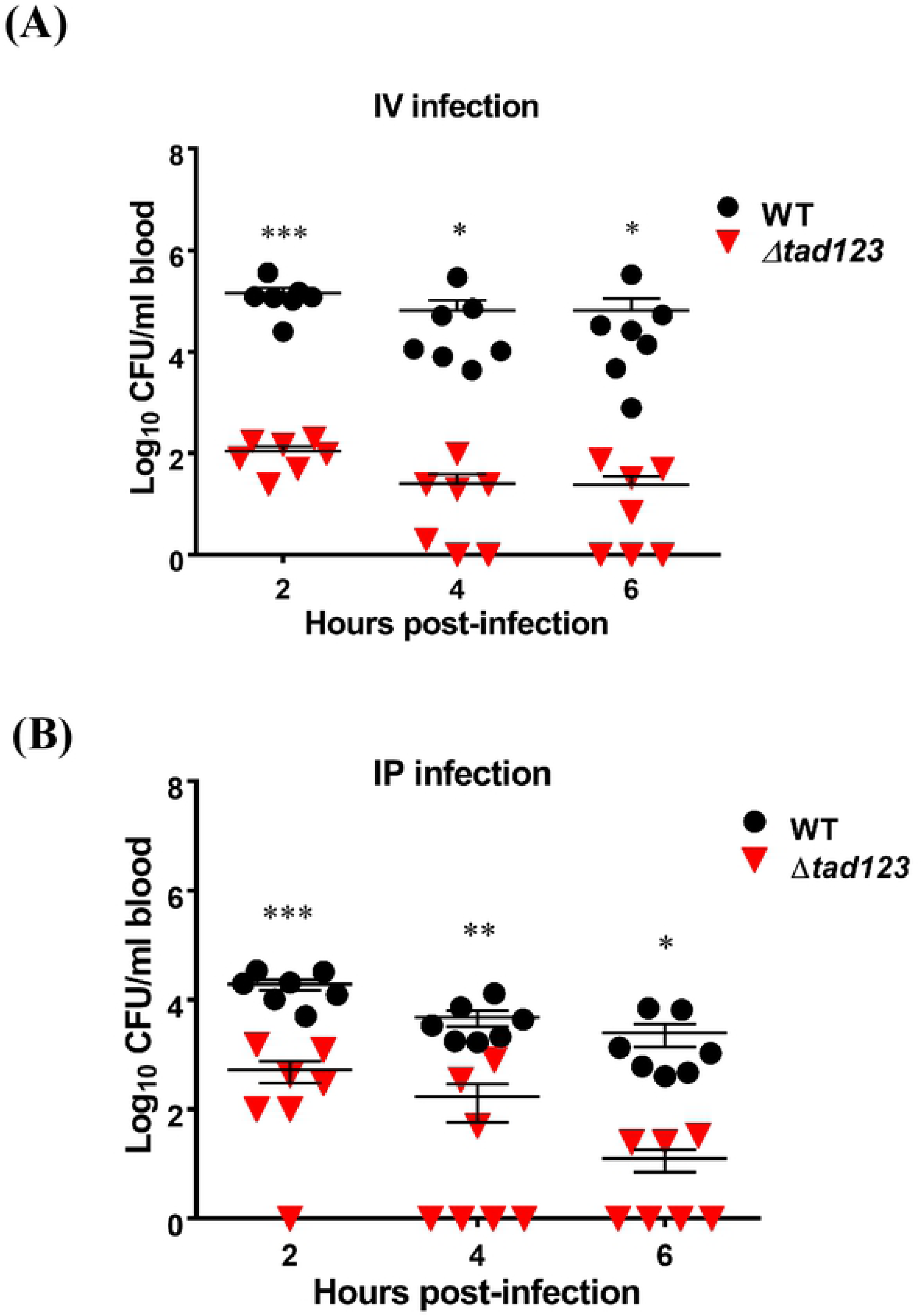
Recovery of *V. vulnificus* from blood after intravenous or intraperitoneal mouse infection. Each mouse was intravenously **(A)** or intraperitoneally **(B)** injected with bacteria that had been previously incubated in the rat peritoneal cavity for 6 hours for *in vivo* adaptation. Blood samples were acquired from infected mice via cardiac puncture at the indicated times. Viable bacterial cells were counted by plating on 2.5% NaCl HI agar plates. Very low numbers of Δ*tad123* mutant cells were recovered from the blood of infected mice. Data shown represent the mean ± SEM of three independent experiments performed in seven mice. Statistical analysis was carried out using Student’s *t* test (*, *P* < 0.05; **, *P* < 0.01; ***, *P* < 0.001).

### Tad pili are required for *V. vulnificus* serum resistance

Serum bactericidal activity is an important innate immune defense against intravascular invasion by bacterial pathogens (31, 32). Thus, we hypothesized that Tad pili might play a protective role against serum components. To address this hypothesis, we tested the susceptibility of triple mutant cells to normal human serum (NHS). Bacterial viability was assessed after 2 hours of incubation with different NHS concentrations. Notably, the bacteria lacking all three *tad* loci were extremely sensitive to human serum (Fig. 8A) (*P* < 0.001). For the Δ*tad123* mutant, 20% NHS led to dramatically decreased viability, and exposure to 40% NHS resulted in the death of up to 95% of the cells. In contrast, the wild-type cells could actively grow in 10 to 50% NHS (Fig. 8A). A time course assay with 50% NHS was carried out to further compare the serum resistance levels of the isogenic mutant stain and the wild-type strain. During the first hour of incubation, more than 90% of the cells of both strains lost viability (Fig. 8B). Thereafter, however, the wild-type strain recovered and resumed rapid growth, whereas most of the Δ*tad123* mutant cells were dead. This result explains the significant difference in the survival rates observed between the wild-type and Δ*tad123* mutant strains and further confirms that Tad pili are required for *V. vulnificus* serum resistance.

**Figure 8.**
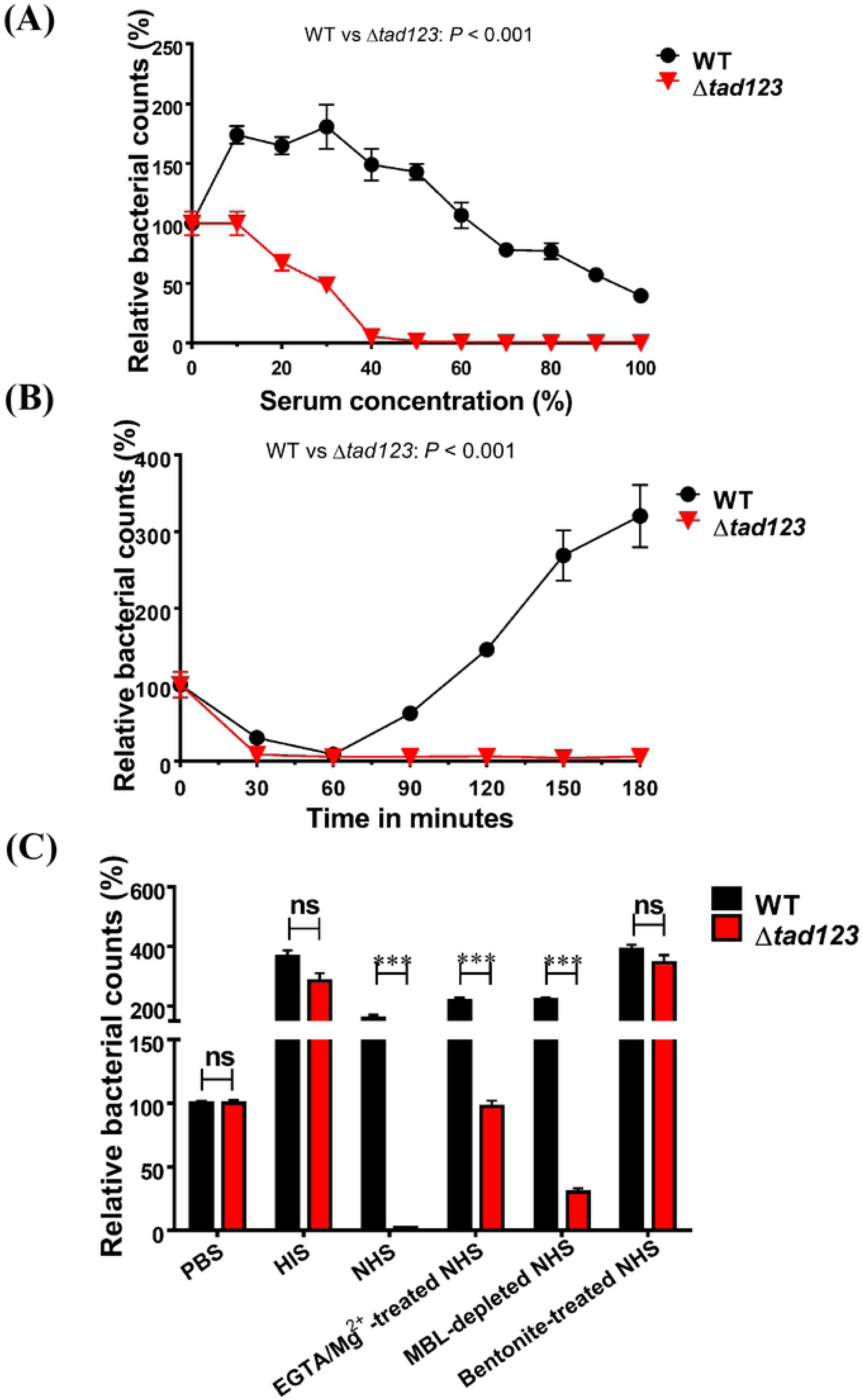
Survival of *V. vulnificus* against human serum bactericidal killing. **(A)** Log-phase bacteria were incubated with different NHS concentrations at 37 °C for 2 h and the numbers of viable cells were determined. **(B)** Log-phase bacteria were incubated with 50% NHS and the numbers of viable cell were determined in a time-dependent manner. **(C)** Bacteria were incubated with HIS, undiluted NHS, EGTA/Mg^2+^-treated NHS, MBL-depleted NHS or bentonite-treated NHS for 2 h. Viable bacterial cells were counted by plating on 2.5% NaCl HI agar plates. Bacteria were incubated in PBS as a control. Data shown represent the mean ± SEM of four independent experiments performed in triplicates. Statistical analysis was carried out using two-way ANOVA followed by the Bonferroni *post hoc* test (A and B) or Student’s *t* test (C). ***, *P* < 0.001; ns, not significant.

### The alternative complement pathway plays a dominant role in the killing of the Δ*tad123* mutant cells

Given that the complement system, which is activated by pathogenic bacteria is primarily responsible for the direct killing of bacteria in NHS (33), we further dissected the bactericidal activity of three complement pathways activated by *V. vulnificus*. Indeed, the use of heat-inactivated serum (HIS) lacking the lytic complement activity successfully rescued the viability of the mutant strain (Fig. 8C). This finding indicates that heat-labile complement proteins are responsible for the killing of the Δ*tad123* mutant cells in NHS. Complement activation occurs via one or more of three pathways: the classical pathway, the MBL/lectin pathway and the alternative pathway (31). To identify which complement pathway was responsible for the death of the Δ*tad123* mutant cells in serum, we selectively blocked the specific complement activation pathways. Remarkably, inhibition of the alternative pathway completely ablated the complement-mediated killing activity. The survival of the Δ*tad123* mutant cells fully recovered to the wild-type level in bentonite-absorbed NHS (Fig. 8C). Furthermore, inhibition of either the classical pathway or the lectin pathway partially recovered the survival of the mutant cells (Fig. 8C) (*P* < 0.001). Taken together, these results indicate that Tad pili likely play an important role in protecting *V. vulnificus* from direct complement-mediated bacteriolysis resulting predominantly from activation of the alternative pathway.

## Discussion

Host-pathogen interactions during microbial infections can be described as a dynamic battlefield where the microbe’s clever strategies for survival and multiplication confront the formidable host immune defenses. To investigate the virulence regulation of *V. vulnificus* during infection, we recently performed comparative genome-wide transcriptional analyses of cells grown *in vitro* and *in vivo.* A rat peritoneal infection model was used to simulate the physiological host milieu. Interestingly, among the newly identified *in vivo*-expressed candidate genes, *tad1* was found to be highly upregulated *in vivo* (unpublished data). The pathogenic potential of the *tad1* cluster is also supported by previous reports of the ubiquity of the *tad1* locus in sequenced virulent *V. vulnificus* strains (14-16). Furthermore, it is notable that the genome of *V. vulnificus* CMCP6 contains three distinct *tad* loci, in which similar functional genes are organized in the same order and transcriptional orientation. In this study, our goals were to investigate why *V. vulnificus* CMCP6 has maintained three *tad* loci throughout evolution and how each *tad* operon contributes to *V. vulnificus* virulence and to determine whether all three *tad* operons are required for its virulence. By deleting each *tad* locus and complementing the deletion *in trans*, we attempted to address these questions (at least in part) and found that all three *tad* operons are required for the full virulence of *V. vulnificus*. Only complete abrogation of all three *tad* loci led to significantly decreased lethality in mice (Table 1). Based on animal and cell culture infection models coupled with molecular genetic analyses, we came to understand the coordinated contributions of the three *V. vulnificus tad* operons to host cell invasion as well as to survival of complement-mediated bacteriolysis. Deletion of all three *tad* loci impaired the adherence of the bacterial cells to the host cells (Fig. 2), thus hampering RtxA1 secretion and cytotoxin delivery (Fig. 5) and, consequently, tissue invasion (Fig. 6). These results corroborate our previous findings that host cell contact is required for *V. vulnificus* toxin secretion and pathogenicity (20).

The bactericidal action of serum is an important component of the host defense against bloodstream infections (31, 32). As most fatal cases of *V. vulnificus* infection result from septicemia, serum resistance is considered an essential feature for survival in the host environment. It is well documented that clinical *V. vulnificus* isolates have a significantly greater survival ability in human serum compared with that of environmental isolates (34, 35). Several mechanisms have been proposed to explain this phenomenon, the most significant of which may be differences in siderophore expression and/or capsule formation (34). In the present study, we discovered a novel function of *V. vulnificus* Tad pili in conferring resistance to the complement-mediated bactericidal activity of its host. The ubiquity of the *tad1* cluster in virulent *V. vulnificus* strains suggests that the surface expression of Tad pili may be another key determinant for the survival of *V. vulnificus* in host milieus, which could conceivably differentiate the clinical and environmental strains. Bacteria lacking Tad pili rapidly lost viability in serum via direct complement-mediated bacteriolysis, predominantly activated via the alternative pathway. The mechanism through which Tad pili protect bacteria from complement attack should be further studied. The poor immunogenicity of Flp pili could have something to do with the serum resistance. Given that the *tad* triple mutant lost its slime-like surface morphology (which could be complemented *in trans* by a cosmid library), it is plausible that Flp pili could anchor secreted polysaccharides during formation of durable capsular lattice.

Inhibition of the alternative complement pathway by the Flp pili might be related to the low immunogenicity of the structural pilin protein. To understand the poor recognition of Tad pili by the immune system, we analyzed the antigenicity and structural characteristics of Tad pilin using bioinformatics tools. After *in silico* prediction of the 3D structure of Tad pilin, we compared it with the orthologs from *Aggregatibacter actinomycetemcomitans*, which seem to have high functional similarity with that of *V. vulnificus. Bordetella pertussis* Fim2 and *Escherichia coli* CfaB (S6 Fig). *V. vulnificus* Tad pilin was predicted to form an alpha helix that partially overlaps structurally with the *A. actinomycetemcomitans* Flp-1 and Flp-2 and *B. pertussis* Fim2 pilins, which are thought to contribute to the assembly of pilin monomers into the fimbrial ultrastructure. When compared with the immunogenic Fim2 and CfaB pilins, which function as vaccine candidates, Tad pilin appeared to be relatively hydrophobic, and only a small fraction contained the hydrophilicity required for antigenicity (S7 Fig).

The presence of pilus-like structures has been reported to be more closely associated with clinical isolates of *V. vulnificus* than with environmental strains (36). Under SEM, we observed filamentous surface structures that extended from the wild-type cell bodies that were absent from the Δ*tad123* triple operon mutant cells (Fig. 3). Interestingly, the presumable Tad pili structure became more elongated when the cells were grown *in vivo*, suggesting a pathogenic function during establishment of successful infections. This morphological change under *in vivo* culture conditions corroborates the previously suggested hypothesis that the Tad pili significantly contribute to *V. vulnificus* pathogenicity (12). The thickness and size of the putative pili structures were quite elusive, unlike other Tad/Flp pili such as those of *Aggregatibacter* (previously *Actinobacillus*) (6-9). To confirm the cell surface expression of Tad pili, we performed immunogold-labeling and fluorescence staining. To the best of our knowledge, no specific ultrastructural analysis of *V. vulnificus* Tad pili and their molecular pathogenic roles have been previously reported. While performing these experiments, we failed to raise functional antibodies against the *V. vulnificus* Flp pilin, presumably because of its very low hydrophilicity and immunogenicity as addressed above. To overcome this obstacle, we first fused a V5 tag sequence to the N- or C-terminus of the *flp-1* gene on the chromosome. However, we could not detect any signal from chromosomally V5-tagged Flp in a dot blot analysis using an anti-V5 antibody (data not shown). The V5-tagged Flp might have been structurally defective, leading to ineffective assembly into pili structures. Alternatively, the engineered strains may have incurred polar effects on downstream gene expression during double crossover homologous recombination. We subsequently constructed a V5-tagged Flp overexpression system encoded on the multicopy pBAD24 plasmid (S4 Fig). Our hypothesis was that when expressed under the control of an arabinose-inducible promoter, some proportion of the overexpressed V5-tagged Flp proteins might be randomly incorporated during assembly of the chromosomally-expressed native pilin subunits. As expected, we could detect positive signals from *V. vulnificus* cells overexpressing the Flp-V5 fusion protein in both experiments (Fig. 4). However, only a fraction of the transconjugants carrying the overexpression plasmid could be stained with immunogold or fluorescence approaches, suggesting minimal incorporation of V5-tagged Flp expressed from the single-copy chromosomal locus possibly due to structural deformations by addition of the V5 tag.

Taken together, our results provide new insights into the pathogenic significance of Tad pili in *V. vulnificus* CMCP6. The Tad provide pathogenic *V. vulnificus* with the ability to adhere to and invade host cells and shield the cells against complement-mediated bacteriolysis inside the host. During these two distinct stages of infection, the Tad pili-mediated host cell adhesion and evasion of the anti-complement activity inside host provide, respectively, the signal required to induce expression of the potent RtxA1 cytotoxin and the ability of *V. vulnificus* to robustly grow *in vivo*.

## Materials and methods

### Bacterial strains, plasmids and media

Bacterial strains and plasmids used in this study are listed in Table 2. *V. vulnificus* CMCP6 is a highly virulent clinical isolate from the Chonnam National University Hospital, South Korea(5, 20). *V. vulnificus* and *E. coli* were grown in 2.5% NaCl heart infusion (HI) and in Luria-Bertani (LB) medium, respectively. Antibiotics were used as previously described (37).

**Table 2.** Strains and plasmids used in this study.

| Strain or plasmid | Description | Source or reference |
| --- | --- | --- |
| Strains |  |  |
| *V. vulnificus* |  |  |
| CMCP6 | Wild-type, clinical isolate | CNU Hospital |
| $\Delta tad1$ | CMCP6 with in-frame deletion of entire structural genes in *tad1* locus | This study |
| $\Delta tad2$ | CMCP6 with in-frame deletion of entire structural genes in *tad2* locus | This study |
| $\Delta tad3$ | CMCP6 with in-frame deletion of entire structural genes in *tad3* locus | This study |
| $\Delta tad12$ | CMCP6 with in-frame double deletion of entire structural genes in *tad12* loci | This study |
| $\Delta tad13$ | CMCP6 with in-frame double deletion of entire structural genes in *tad13* loci | This study |
| $\Delta tad23$ | CMCP6 with in-frame double deletion of entire structural genes in *tad23* loci | This study |
| $\Delta tad123$ | CMCP6 with in-frame triple deletion of entire structural genes in *tad123* loci | This study |
| *V<sub>v</sub>*-pBAD | CMCP6 carrying pBAD24 empty vector | This study |
| *V<sub>v</sub>*-FlpV5 | CMCP6 carrying pCMM2103 | This study |
| $\Delta tad$-FlpV5 | $\Delta tad123$ carrying pCMM2103 | |
| *E. coli* |  |  |
| BL21 (DE3) | F<sup>−</sup> *hsdS*(r<sub>B</sub><sup>−</sup> m<sub>K</sub><sup>−</sup>) *gal* with T7 RNA polymerase gene in chromosome | Novagen |
| DH5α | F<sup>−</sup> *recA1* restriction negative | Laboratory collection |
| SY327 $\lambda$ *pir* | $\Delta$(*lac pro*) *argE* (Am) *rif<sup>r</sup>nalA recA56* $\lambda$ *pir* lysogen | (38) |
| SM10 $\lambda$ *pir* | *thi thr leu tonA lacY supE recA::RP4-2-Tc<sup>r</sup>::Mu Km<sup>r</sup>* $\lambda$ *pir* lysogen | (38) |
| Plasmids |  |  |
| pBAD24 | Expression vector, arabinose inducible promoter; Amp<sup>r</sup> | (48) |
| pET30a+ | Km<sup>r</sup> *E. coli* cloning vector with T7 promoter upstream of N-terminal His<sub>6</sub> tag | Novagen |
| pDM4 | Suicide vector with *oriR6K sacB*; Cm<sup>r</sup> |  |
| pLAFR3 | IncP cosmid vector; Tc<sup>r</sup> | (49) |
| pRK2013 | IncP Km<sup>r</sup> Tra Rk2<sup>+</sup> *repRK2 repE1* | (50) |
| pCMM2103 | pBAD24 expressing a fusion protein of Flp-1 pilin and V5 tag | This study |
| pCMM2104 | pDM4 with a 2-kb SmaI/XhoI in-frame deleted *tad1* operon | This study |
| pCMM2105 | pDM4 with a 2-kb SmaI/XhoI in-frame deleted *tad2* operon | This study |
| pCMM2106 | pDM4 with a 2-kb SmaI/XhoI in-frame deleted *tad3* operon | This study |
| pCMM2107 | pLAFR3 with a 25-kb fragment containing the in-frame *tad1* operon | This study |
| pCMM2108 | pLAFR3 with a 25-kb fragment containing the in-frame *tad2* operon | This study |
| pCMM2109 | pLAFR3 with a 25-kb fragment containing the in-frame *tad3* operon | This study |

Cm<sup>r</sup>, Cm resistance; Tc<sup>r</sup>, Tc resistance; Amp<sup>r</sup>, Amp resistance; Km<sup>r</sup>, Km resistance.

### Locus deletion mutant construction and complementation

We constructed in-frame single, double, and triple deletion mutants of entire structural genes in the *tad1, tad2* and *tad3* loci by the allelic-exchange method (38). We designed two sets of primers to amplify ~1-kb DNA fragments in the upstream or downstream region of each *tad* operon (S1 Table). The primers were synthesized with overhangs recognized by specific restriction enzymes (REs). The upstream and downstream amplicons of each *tad* operon were ligated by cross-over PCR to produce a 2-kb fragment (39). The fusion fragments were digested with appropriate REs and subcloned into pDM4 suicide vector. The resulting recombinant vector was transformed into *E. coli* SM10 λ *pir* and subsequently transferred into *V. vulnificus* CMCP6 by conjugation. Stable Cm^R^ transconjugants were selected on *Vibrio*-selective thiosulfate citrate bile salt sucrose (TCBS) agar plate containing Cm. Plating of the transconjugants on 2.5% NaCl HI agar plate containing 10% sucrose was performed to select clones that experienced second homologous recombination events forcing excision of the vector sequence and leaving only mutated or wild-type allele of the genes. Each in-frame deletion mutation was confirmed by PCR with the chromosomal DNA from the respective mutant as template.

For the use in genetic complementation experiments, we screened cosmid clones that contain intact *tad1, tad2* or *tad3* operon from a pLAFR3 cosmid library of *V. vulnificus* CMCP6 (20, 40-42). The selected cosmid library clone was transferred to the triple *tad* operon deletion mutant by triparental mating with a conjugative helper plasmid pRK2013. The transconjugants were screened on TCBS agar plates containing tetracycline and confirmed by PCR. To fulfill Koch’s postulates, we performed a complementation analysis. The Δ*tad123* mutant was separately complemented with an individual cosmid clone harboring each *tad* operon. The restoration of *tad* operon expression was confirmed by the conventional RT-PCR (S3 Fig).

### Conventional and real-time RT-PCR

The transcriptional levels of the three structural *flp* genes, which encoded the major structural components of Flp pili, were measured by conventional and real-time RT-PCR. *gyrA* was chosen as the reference gene forn the qRT-PCR as previously reported (43). Forward and reverse primer pairs were designed and are provided in S2 Table. Total RNA was isolated from log-phase bacterial cells grown in the rat peritoneal cavity or in 2.5% NaCl HI broth using the RNeasy minikit (Qiagen). One microgram of purified RNA was converted into cDNA using QuantiTect® Reverse Transcription Kit (Qiagen) in accordance with the manufacturer’s protocol. qRT-PCR was performed to quantify each target transcript using QuantiTect SYBR green PCR kit (Qiagen). The relative gene expression was normalized to the expression of *gyrA* using the threshold cycle (ΔΔC_T_) method (44). For conventional RT-PCR, 16S rRNA was used as the internal standard. After 25 to 35 cycles, the amplicons were separated on 2% (wt/vol) agarose gels and stained with ethidium bromide. The transcription levels of *flp-1* under iron-limited and solid surface growth conditions were also analyzed by qRT-PCR. For the iron limitation experiment, dipyridyl (Sigma-Aldrich) was added to the 2.5% NaCl HI broth at a final concentration of 80 µM for iron limitation.

### Ethics Statement

All animal experimental procedures were performed with approval from the Chonnam National University Institutional Animal Care and Use Committee under protocol (H-2015-44) and were conducted in accordance with the guidelines of the Animal Care and Use Committee of the Chonnam National University.

### LD_50_ determination

The intraperitoneal 50% lethal dose (i.p. LD_50_) of *V. vulnificus* was determined using 7-week-old, randomly bred specific-pathogen-free (SPF) female ICR mice (Daehan Animal Co., Daejeon, South Korea). Five mice per group were intraperitoneally inoculated with 10-fold serial dilutions of fresh bacterial suspensions (10^9^ to 10^5^ CFU/mouse). The intragastric (i.g.) LD_50_ was determined using six-day-old randomly bred SPF CD-1 suckling mice (Daehan Animal Co., Daejeon, South Korea). Seven mice per group were intragastrically administered with 10-fold serial dilutions of fresh bacterial suspensions containing 0.1% Evans blue (Sigma-Aldrich) to ensure correct i.g. administration. The control animals received 100 µl of PBS containing 0.1% Evans Blue. The challenged mice were monitored for 48 h. LD_50_ values were calculated based on probit analysis, using IBM SPSS 21.0 software (IBM).

### Construction of C-terminally V5-tagged Flp (Flp-V5)

DNA fragments of the structural *flp-1* pilus gene without its stop codon were amplified and subcloned into the pBAD24 vector. Subsequently, double-stranded oligonucleotides encoding the V5 peptide, “GGTAAGCCTATCCCTAACCCTCTCCTCGGTCTCGATTCTACGTAA”, were fused to the C-terminus of the *flp-1* gene in the pBAD24-Flp plasmid. At the end of the V5 sequence, a TAA codon was added to terminate translation. The pBAD24 plasmids containing the Flp-V5 fusion protein were transformed into *E. coli* DH5α competent cell. The sequence of the cloned fragment was confirmed by DNA sequencing. The resulting vectors were transferred into *V. vulnificus* via triparental mating with a conjugative helper plasmid pRK2013. The transconjugants were screened on 2.5% NaCl HI agar plates containing ampicillin.

### Dot blot analysis

*V. vulnificus* strains carrying pCMM2103 (pBAD24::Flp-V5) were grown for 4 h on 2.5% NaCl HI agar plates containing ampicillin. Flp-V5 expression was induced for 4 h via addition of 0.1% L-arabinose. The cell suspensions were applied to nitrocellulose membrane and fixed with 4% paraformaldehyde for 20 min. The membrane was blocked for 1 h using 5% skim milk in PBS and then incubated with anti-V5 polyclonal antibodies (diluted 1:5000, Abcam) for 2 h. After washing, the membrane was developed with HRP-conjugated goat anti-rabbit IgG secondary antibody (Dako). Stained dots on a white background indicated positive results.

### Scanning electron microscopy

To compare surface structure of the wild-type, Δ*tad123* and Δ*tad123* (pLAFR3::*tad1*), we performed SEM observation. Bacteria were grown *in vivo* using a rat peritoneal infection model as previously described (41). To minimize shearing force during bacterial preparations, all procedures were carefully performed. And we also applied osmotic adaptation with fixative by 3 staged applying a step-down approach from 2.5% NaCl containing fixative to 0.9% and 0% NaCl containing solution with a gentle agitation. Bacterial cells were fixed at room temperature for 4 h in a fixation solution containing 0.5% glutaraldehyde and 4% paraformaldehyde in 0.05 M sodium cacodylate buffer (pH 7.2). After three washes with 0.05 M cacodylate buffer, all of the samples were mounted on nickel grids coated with carbon film (150 mesh) (EMS, USA). After blocking nonspecific binding sites with 1 % BSA in EM-immunogold (EMG) buffer (0.05% Tween, 0.5 M NaCl, 0.01 M phosphate buffer, pH 7.2), the samples were incubated at 4°C for 24 h with anti-V5 tag monoclonal antibody (ab27671, Abcam, UK) at a 1:20 dilution at 4°C, followed with incubation for 1 h in goat anti-rat antibody (1:50) conjugated to 6 nm gold particles. The grids were washed in EMG buffer, PBS, and distilled water and stained for 12 min with 4% uranyl acetate in deionized distilled water. The surfaces of all of the samples were observed using a field emission scanning electron microscope (Helios G3 CX, FEI Co., Hillsboro, Oregon, USA) at 1 kV acceleration with TED mode.

### Immunogold labellng

A drop of *V. vulnificus* cell suspension was applied to a nickel cell suspension was applied to a nickel grid coated with carbon film for 1 min. Because of its structural fragility of *V. vulnificus*, we prepared the bacterial sample with very gentle manner such as limited frequencies of pipetting and washing processes. Moreover, to minimize the insults from the critical points drying, we firstly fixed the *in vivo* grown cells with osmolarity-modified fixative (which contains 2.5% NaCl) and changed the solution to conventional fixative for SEM study (0.5% glutaraldehyde and 4% paraformaldehyde in 0.05 M sodium cacodylate buffer (pH 7.2)) under room temperature. Moreover, to reduce damages from electron and enhance the resolution beam during SEM analysis, we used the focused ion scanning electron microscope (FIB). Subsequently, the samples were incubated with the anti-V5 polyclonal antibody (diluted 1:20, Abcam) and labeled with 5-nm colloidal gold-conjugated goat anti-rabbit IgG secondary antibody (diluted 1:20, BritishBioCell, UK).

### Transmission electron microscopy

*V. vulnificus* cells were fixed in a fixation solution containing 0.5% glutaraldehyde and 4% paraformaldehyde in 0.05 M sodium cacodylate buffer (pH 7.2) at room temperature for 4 h. After three washes with 0.05 M cacodylate buffer, all of the samples were mounted on nickel grids coated with carbon film (150 mesh) (EMS, USA). After staining with 2% uranyl acetate, the samples were examined with a transmission electron microscope (TEM) (JEM-1400; JEOL Ltd., Japan) at 80 kV acceleration.

### Confocal microscopy

To induce V5-tagged pilin expression, mid-log phase *V. vulnificus* cells were grown for 4 h on 2.5% NaCl HI-ampicillin agar plates supplemented with 0.1% L-arabinose. Bacterial pellet was then gently suspended in PBS buffer. The induced bacterial cells were directly immobilized on poly-L-lysine-coated coverslips. Samples were fixed with 4% formaldehyde for 30 min and then incubated with primary anti-V5 antibodies (1:300) for 2 h. After three washes with PBS buffer, the cells were incubated for 1 h with a Texas Red-conjugated anti-rabbit secondary antibody (Molecular Probes) and DAPI (Invitrogen). The samples were observed under a laser scanning confocal microscope (LSM 510, Zeiss, Oberkochen, Germany), and the obtained images were analyzed by using the ZEN Lite software (Zeiss, Oberkochen, Germany).

### Adhesion assay

HeLa cell (Korean Cell Line bank) monolayers were grown on chamber slides (Nunc) and then infected for 45 min with log-phase *V. vulnificus* strains at an MOI of 250. Longer incubation times were tested to determine the adhesive recovery of the Δ*tad123* mutant. The monolayer was washed twice with PBS to remove non-adherent bacteria. Following the last wash, the cells were fixed in methanol and stained with 0.1% Giemsa (Sigma-Aldrich). The number of *V. vulnificus* cells that adhered to single HeLa cells was enumerated and examined under a light microscope at 400× and 1,000× magnification (Nikon Eclipse 50i, Japan).

### Western blot analysis

To detect RtxA1, HeLa cells grown in 6-well plates were infected for 35 and 45 min with log-phase *V. vulnificus* strains at an MOI of 100. The bacteria attached to the HeLa cells were lysed using a lysis buffer (Cell Signaling), followed by concentration using an Amicon Ultra-0.5 centrifugal filter apparatus (Merck KgaA). The samples were then subjected to 10% SDS-PAGE. RtxA1 proteins were detected using an anti-GD domain antibody (RtxA1-C, a band of approximately 130 kDa) (25).

### Cytotoxicity assay

To determine the effect of *tad* operon mutations on cytotoxicity against HeLa cells, we performed the lactate dehydrogenase (LDH) release assay as previously described (37).

### *In vivo* invasion assay

Bacterial cells that translocated from the intestine to the bloodstream were measured as previously described (43). Seven-week-old randomly bred SPF female ICR mice were starved for 16 h. The ileum was tied off in a 5-cm segment and log-phase *V. vulnificus* cells (4.0 × 10^6^ CFU/400 µl) were inoculated into the ligated segment. Blood samples were acquired from the infected mice via cardiac puncture. The number of viable bacterial cells was counted by plating on 2.5% NaCl HI agar plates. In parallel, viable *V. vulnificus* cells in the ligated ileal loops were also enumerated by plating on TCBS agar plates.

### *In vitro* invasion assay

Polarized HCA-7 cells (from Professor Eayl Raz, University of California San Diego) grown in Transwell^®^ filter chambers (8 μm pore size; CoStar, Cambridge, MA, USA) were apically exposed to log-phase *V. vulnificus* cells at an MOI of 5. Invasiveness was determined by measuring the number of bacterial cells that translocated from the apical to basolateral compartment of the Transwells. Viable bacterial cells were counted by plating on 2.5% NaCl HI agar plates.

### Determination of *in vivo V. vulnificus* growth

*In vivo* growth of *V. vulnificus* was measured using the dialysis tube implantation model as previously described (43). CelluSep^®^ H1 dialysis tubing (MWCO 12,000~14,000; Membrane Filtration Products, Inc. Texas) was incubated with PBS overnight. The dialysis tube was disinfected with 70% alcohol for 1 h and washed three times with sterile PBS before use. Seven-week-old female Sprague Dawley (SD) rats (DBL. Co. Ltd, Daejeon, Korea) were anesthetized with a mixture of 10% Zoletil and 5% Rumpun dissolved in PBS. Three 10-cm dialysis tubes containing 2 ml of 5 × 10^5^ CFU/ml *V. vulnificus* cells were surgically implanted into the rat peritoneal cavity. The bacterial growth at each time point was analyzed using three rats. Culture samples were harvested for viable cell counting on 2.5% NaCl HI agar plates 2, 4 and 6 h after implantation.

### *V. vulnificus* growth in mouse blood

To assess bacterial growth in blood, seven mice per group were intravenously (i.v.) or i.p. injected with 100 µl of 5 × 10^5^ CFU cells that had been incubated in the rat peritoneal cavity for 6 h for *in vivo* adaptation. Blood samples were acquired from the infected mice via cardiac puncture at the indicated times. Viable bacterial cells were counted by plating on 2.5% NaCl HI agar plates.

### Determination of serum bactericidal activity against *V. vulnificus*

Log-phase *V. vulnificus* cells (1.0 × 10^7^ CFU/10 µl) were added to 200 µl of PBS containing various NHS concentrations. The samples were incubated at 37°C for 2 h. Viable bacterial cells were counted by plating on 2.5% NaCl HI agar plates. To block activation of the classical pathway, NHS was pretreated with 10 mM ethylene glycol-bis (2-aminoethylether)-*N, N, N*′, *N*′-tetraacetic acid and 5 mM MgCl_2_ for 30 min at 37°C (EGTA/Mg^2+^) (45). To prepare the MBL-depleted serum, mannose-agarose beads (Sigma-Aldrich) were washed three times with sterile PBS and then incubated with NHS at 4°C for 1 h with gentle rotation (46). The alternative pathway is inhibited via properdin absorption with bentonite (47). 10 mg of bentonite was washed three times with PBS and incubated with NHS at 37°C for 10 min to absorb the properdin.

### Statistical analysis

The results are expressed as the mean ± standard error of the mean (SEM) unless otherwise stated. Each experiment was repeated a minimum of three times, and the results from representative experiments are shown. Statistical analyses were performed using the Prism 5.00 software for Windows (GraphPad software, San Diego, CA). Multiple comparisons were performed using Student’s *t* test and analysis of variance (ANOVA) followed by Bonferroni *post hoc* tests. *P*-values < 0.05 was considered statistically significant.

## Supporting Information Legends

**S1 Fig. *Vibrio vulnificus* CMCP6 *tad* loci.**

**S2 Fig. Significantly prolonged survival of mice intraperitoneally infected with the Δ*tad123* mutant strain (n=5).** Seven-week-old randomly bred SPF female ICR mice were intraperitoneally infected with 1 x10^7^ CFU/mouse **(A)** or 1 x10^6^ CFU/mouse **(B)** of fresh bacterial suspensions. The challenged mice were monitored for 48 h. Statistical analysis was carried out using Kaplan-Meier analysis followed by the log-rank test (**, *P* < 0.01).

**S3 Fig. Restoration of structural *flp* gene expression in *tad* operon**-**complemented strains.** Expression of the structural *flp* genes was assessed using conventional RT-PCR. RNA was isolated from log-phase bacteria grown in 2.5% NaCl HI broth and then converted into cDNA. RT-PCR was performed using primers specific for each structural *flp* gene as shown in Table S2. The 16S *rRNA* housekeeping gene was employed as the internal control.

**S4 Fig. Detection of V5-tagged Flp fusion proteins from induced *E. coli* cells by Western blot analysis.** Bacteria were grown in LB Amp broth supplemented with (inducing) or without (non-inducing) 0.1% arabinose for 4 h. The Flp-V5 fusion proteins were detected using an anti-V5 polyclonal antibody.

**S5 Fig. Growth of *V. vulnificus* in high-glucose DMEM.** Log-phase *V. vulnificus* cells were grown in high-glucose DMEM, and the OD_600_ was measured every two hours for 8 h. The growth pattern of the Δ*tad123* mutant cells was identical to that of the wild-type strain. Data shown represent the mean ± SEM of three independent experiments performed in triplicate.

**S6 Fig. Comparison of *V. vulnificus tad* with other bacterial pilins.** The predicted 3D structure of *V. vulnificus* Tad pilin (hot pink) was overlaid with those of *A. actinomycetemcomitans* Flp1 **(A, yellow)** and Flp2 **(B, orange)**, *B. pertussis* Fim2 **(C, cyan)**, and *E. coli* CfaB **(D, green)**.

**S7 Fig. Hydrophilicity comparison of *V. vulnificus* Tad pilin (red) with immunogenic *B. pertussis* Fim2 (blue) and *E. coli* CfaB (green).** Positive values indicate hydrophilicity while negative values indicate hydrophobicity. The red line shows the average hydrophilicity scores of the Fim2 and CfaB antigenic domains. A minor fraction adjacent to the alpha helical region of Tad pilin possesses less than 0.5 hydrophilicity.

**S1 Table. Primers used for the construction of *tad* operon deletion mutants and the fusion protein.**

**S2 Table. Primers used in the RT-PCR study**

## Acknowledgements

J.H.R. was supported by the National Research Foundation of Korea (NRF) grant funded by the Korea government (MSIT) (No. 2018R1A5A2024181), by the Bio & Medical Technology Development Program of the NRF funded by the Korean government, MSIP (NRF-2017M3A9E2056372) and by a grant from the National Program for Cancer Control, Ministry of Health & Welfare, Republic of Korea A17C0038(1720120). S.E.L. was supported by an NRF grant from the MSIP (NRF-2016R1A2B4009611) of the Republic of Korea.

## Author Contributions

Substantial contributions to conception and design of this study: T-MD, KJ, LSE, RJH;

Acquisition of data: T-MD, KJ, SYK, WT, PS, KHL;

Analysis and interpretation of data: T-MD, KJ, LSE, RJH;

Drafting the article: T-MD, KJ, LSE, RJH;

Revising it critically for important intellectual content: T-MD, KJ, LSE, RJH;

Final approval of the version to be published: LSE, RJH

## Declaration of Interests

The authors declare no competing interests.

